# Visual capture of the head-direction signal in a rat model of Fragile X Syndrome

**DOI:** 10.64898/2026.11.06.764867

**Authors:** Adrian J. Duszkiewicz, Anna A. Råstedt, Anuj Vadher, Emma R. Wood, Adrien Peyrache, Paul A. Dudchenko

**Affiliations:** School of Psychology, University of Stirling, Stirling, UK; Division of Neuroscience, University of Manchester, Manchester, UK; Institute for Neuroscience and Cardiovascular Research & Simons Initiative for the Developing Brain, University of Edinburgh, Edinburgh, UK; Montreal Neurological Institute and Hospital, McGill University, Montreal, Canada

## Abstract

Fragile X Syndrome (FXS) is a common inherited monogenic cause of autism spectrum disorder and intellectual disability (ASD/ID), conditions frequently associated with altered sensory processing and multisensory integration. However, the circuit mechanisms underlying these alterations remain poorly understood. Here, we investigate the integration of visual and self-motion information in the head-direction (HD) system in a rat model of FXS (*Fmr1*^*-/y*^ rats). Using high-density silicon probes, we recorded neuronal activity in the postsubiculum (PoSub), a cortical hub of the HD system, while adult *Fmr1*^*-/y*^ and wild-type (WT) rats experienced repeated rotations of a visual landmark. In WT rats, the HD signal only partially followed landmark rotations, consistent with weighted integration of visual and self-motion information, and visual landmark control progressively weakened across repeated cue conflicts. In contrast, the HD signal in *Fmr1*^*-/y*^ rats closely followed landmark rotations and failed to show experience-dependent recalibration. This enhanced visual capture was accompanied by reduced angular head velocity tuning in PoSub neurons, indicating weakened representation of self-motion information. Together, these findings reveal an altered balance between visual and self-motion inputs to the HD system in a rat model of FXS and identify a possible circuit-level mechanism through which sensory integration of vision and self-motion may be disrupted in ASD/ID.

## Introduction

Altered higher-order sensory processing is a common feature of autism spectrum disorder and intellectual disability (ASD/ID) (Wallace et al., 2020) with ample evidence for atypical integration of different sensory streams during both perception and action (larocci and McDonald, 2006; Whyatt and Craig, 2013). Although such alterations have been documented across sensory modalities and behavioural domains in both humans (Feldman et al., 2018) and rodent models of ASD/ID (Monday et al., 2023), the underlying circuit mechanisms and computations remain poorly understood. Navigation provides a particularly compelling context in which to study sensory integration, as spatial awareness requires continual evaluation of allothetic information about the external environment as well as idiothetic signals related to one’s own movements (Poulter et al., 2018; Zhao and Warren, 2015). Spatial navigation thus offers a principled behavioural domain in which altered sensory integration can be linked to defined circuit mechanisms in animal models of ASD/ID.

Sensory integration during navigation depends on interacting hippocampal and extra-hippocampal circuits that continuously update an internal estimate of position and orientation (Wiener et al., 2002; McNaughton et al., 2006). Among these, the rodent head-direction (HD) circuit, the ‘neural compass’, stands out as a highly tractable access point to this integrative function of the cognitive map (Knierim et al., 1995; Yoganarasimha and Knierim, 2005). Neural activity in the HD circuit encodes a circular latent variable, the ‘HD signal’, that precisely maps onto the animal’s current heading. To maintain this accurate representation, the HD signal must be anchored to the allocentric reference frame. The HD circuit achieves this via integration of self-motion signals like the angular head velocity (AHV) with landmark-based inputs (Taube, 2007; Yoder et al., 2011b, 2011a). Substantial experimental and theoretical work suggests that when these two sensory streams are in conflict, they are not combined in a winner-take-all manner but instead through a Bayesian-like weighted integration, with the weights modified by experience (Zugaro et al., 2000; Knight et al., 2014; Jeffery et al., 2016; Page and Jeffery, 2018). When a cue conflict is detected, the HD signal settles at an intermediate anchoring state between the states predicted by landmark or self-motion inputs alone (Zugaro et al., 2000; Knight et al., 2014). Such integration is likely to involve the postsubiculum (PoSub), a cortical hub of the HD circuit (Duszkiewicz et al., 2025) where visual (Siegenthaler et al., 2025) and self-motion (Spalla et al., 2022) information converges to allow for sensory integration (Goodridge and Taube, 1997).

Here, we show that in a rat model of Fragile X syndrome (FXS), a monogenic inherited cause of ASD/ID, the HD signal in PoSub fails to implement the reliability-weighted compromise typically observed in cue conflict conditions. Rather than settling at an intermediate anchoring state, HD tuning in *Fmr1*^*-/y*^ rats is fully captured by vision, consistent with a shift in sensory weighting away from self-motion cues. In line with this, we find reduced AHV tuning in PoSub neurons, indicating weakened cortical representation of idiothetic input that would otherwise constrain reorientation following cue conflict.

## Results

To reliably induce cue conflict over repeated trials within the same session, we used an immersive visual reorientation paradigm (Ajabi et al., 2023). Adult male *Fmr1*^*-/y*^ rats and their wild-type (WT) littermates freely explored an elevated platform within a darkened enclosure, with black curtains minimizing the influence of uncontrolled distal visual cues. The platform was surrounded by a 360° octagonal LED display on which a single salient polarizing visual landmark (a ‘cue’) was presented (Figure 1a). This cue was the only source of illumination in the arena and could be shifted to another position while rats maintained ongoing self-motion, enabling us to quantify the relative contributions of visual landmark input and idiothetic signals during reorientation. Each session began with a baseline epoch in which the cue was fixed at the screen position corresponding to 0°, followed by a sequence of alternating clockwise and counterclockwise cue ‘rotations’ by either 45° or 90°, presented in blocks (16 rotations per session). Rats in both groups readily explored the platform throughout the whole duration of the protocol (Figure S1a) and did not significantly differ in their overall behaviour (Figure S1b-c).

**Figure 1.**
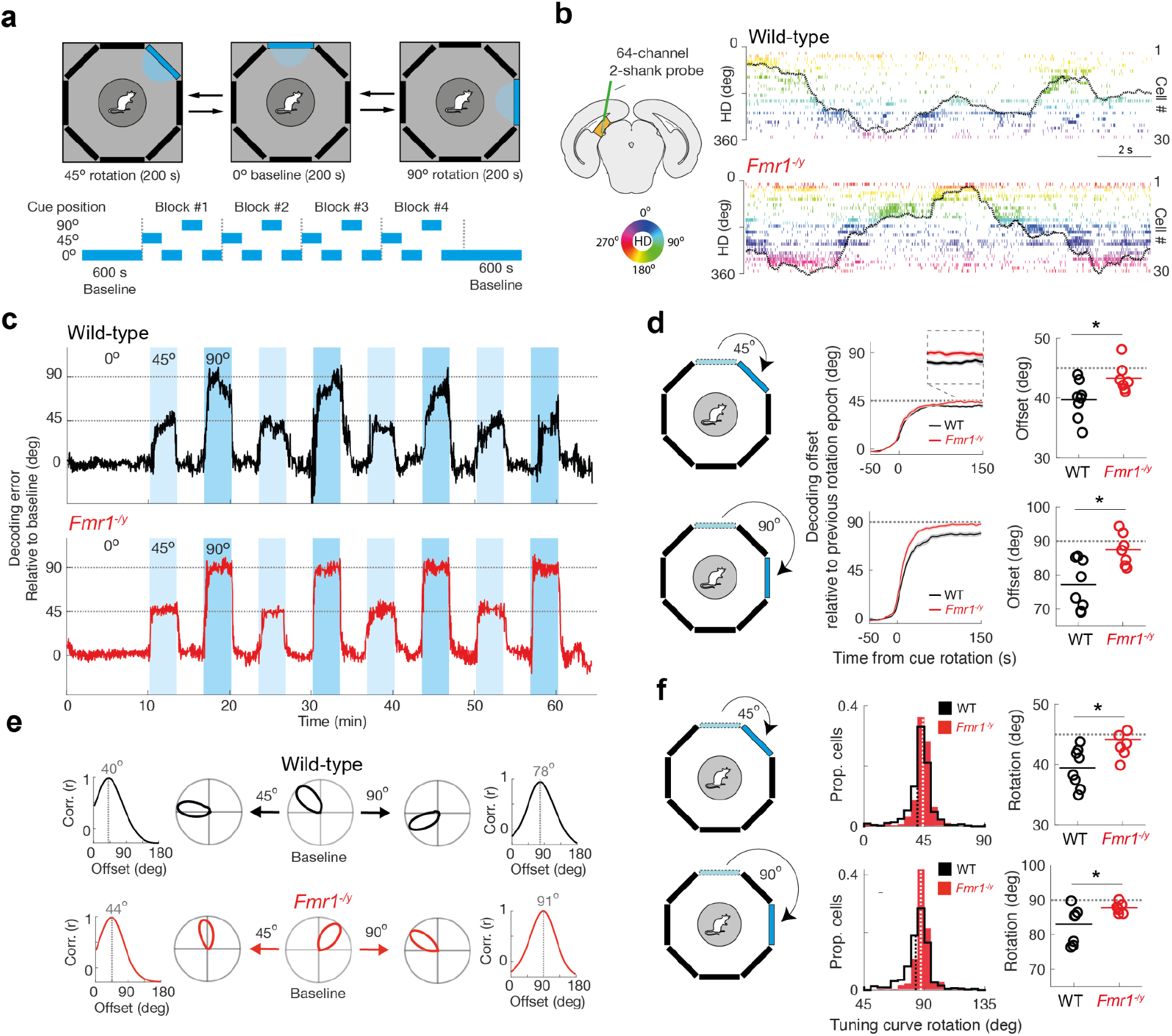
The hallmark of weighted cue integration during HD reorientation is absent in *Fmr1*^*-/y*^ rats. **a**, Visual reorientation apparatus and paradigm. **b**, Silicon probe placement in PoSub (left) and example raster plots of putative PoSub excitatory cells recorded in wild-type rats (top right) and *Fmr1*^*-/y*^ rats (bottom right), with cells sorted according to their preferred direction during the baseline period (30 cells with the highest HD information). Black curves, real HD. **c**, Decoding error plotted for the whole recording session, showing reliable reorientation of the HD signal following individual cue rotations in both wild type (top) and *Fmr1*^*-/y*^ rats (bottom). **d**, Change in the decoding offset of the HD signal around the time of the cue rotation (mean +/-SD) and average decoding offset per animal for 45^°^ rotations (top) and 90^°^ rotations (bottom), showing a tendency for under-rotations in wild-type rats and tendency for accurate rotations in *Fmr1*^*-/y*^ rats (Watson-Williams test on animal averages, 45^°^ rotations: t(13) = 2.61, P < 0.05; 90^°^ rotations: t(13) = 2.54, P < 0.05). **e**, Example HD cell tuning curves in wild type rats (top) and *Fmr1*^*-/y*^ rats (bottom) in the baseline epoch (middle) and following cue rotations (left, right), as well as circular cross-correlations showing under-rotations in wild-type rats and precise rotations in *Fmr1*^*-/y*^ rats. **f**, Individual HD tuning curves tend to under-rotate in WT rats but rotate precisely in *Fmr1*^*-/y*^ rats (t-test on animal averages, 45^°^ rotations: t(13) = 2.22, P < 0.05; 90^°^ rotations: t(13) = 2.30, P < 0.05). Dashed lines on the histograms indicate population medians; horizontal lines on animal average panels are means of animal averages.

To quantify how the HD signal realigns itself in response to conflicting sensory information we recorded populations of neurons in the postsubiculum (PoSub), the postulated site of vestibulo-visual integration in the HD system (Duszkiewicz et al., 2025; Goodridge and Taube, 1997; Yoder et al., 2011b), using silicon probes while rats performed the reorientation task (Figure 1b, Figure S1d-e; 2 sessions per rat; n = 25-97 cells per session from 8 WT rats and n = 35-118 cells per session from 7 *Fmr1*^*-/y*^ rats). This enabled high-precision inference of the internal HD representation from population activity (Figure S1h). We found no differences in the magnitude or stability of HD tuning across the groups (Figure S1f-g).

To assess the magnitude of reorientation, we computed the instantaneous ‘offset’ of the HD signal as the angular mismatch between the population-decoded HD and the animal’s true HD derived from behavioural tracking. Rotations of the visual cue by 45° or 90° reliably evoked reorientation events in both WT and *Fmr1*^*-/y*^ rats, expressed as abrupt increases in offset (Figure 1c). In WT rats, decoded HD often under-rotated its anchoring offset relative to the cue displacement, consistent with reliability-weighted integration of landmark and self-motion information. By contrast, in *Fmr1*^*-/y*^ rats, the offset frequently matched the magnitude of cue rotation closely, indicative of a complete capture by the visual landmark. Accordingly, the mean magnitude of HD reorientation was significantly larger in *Fmr1*^*-/y*^ rats than in WT controls for both 45° and 90° rotations, and did not differ from the cue displacement expected under complete visual capture (Figure 1d). We confirmed this effect at the single-cell level by estimating inter-epoch shifts in HD tuning curves of putative excitatory (EX) cells (n = 334 cells from 8 WT rats, n = 424 cells from 7 *Fmr1*^*-/y*^ rats), computed separately for the 0°, 45°, and 90° epochs (Figure 1e-f). Together, these results indicate that cue-conflict reorientation in *Fmr1*^*-/y*^ rats is biased toward visual landmark control and exhibits reduced evidence of the compromise dynamics characteristic of weighted cue integration.

Since perceived cue reliability is shaped by experience (Goodridge et al., 1998; Knight et al., 2014; Jeffery et al., 2016) we next asked whether the relative weighting of visual landmarks versus self-motion cues evolved across repeated cue-conflict events within a session. Leveraging the multiple reorientation trials per animal, we quantified changes in reorientation magnitude as a function of trial number. For 45° cue rotations, offset magnitude remained stable across repetitions in both WT and *Fmr1*^*-/y*^ rats, with no detectable effect of experience (Figure 2a). In contrast, for 90° rotations WT rats showed a progressive reduction in offset magnitude over successive trials, consistent with experience-dependent down-weighting of the visual landmark and/or increased reliance on self-motion information (Figure 2b). This adaptive reduction was absent in *Fmr1*^*-/y*^ rats, which maintained near-complete visual cue capture across repetitions, indicating impaired experience-dependent recalibration of sensory weights during large cue conflicts.

**Figure 2.**
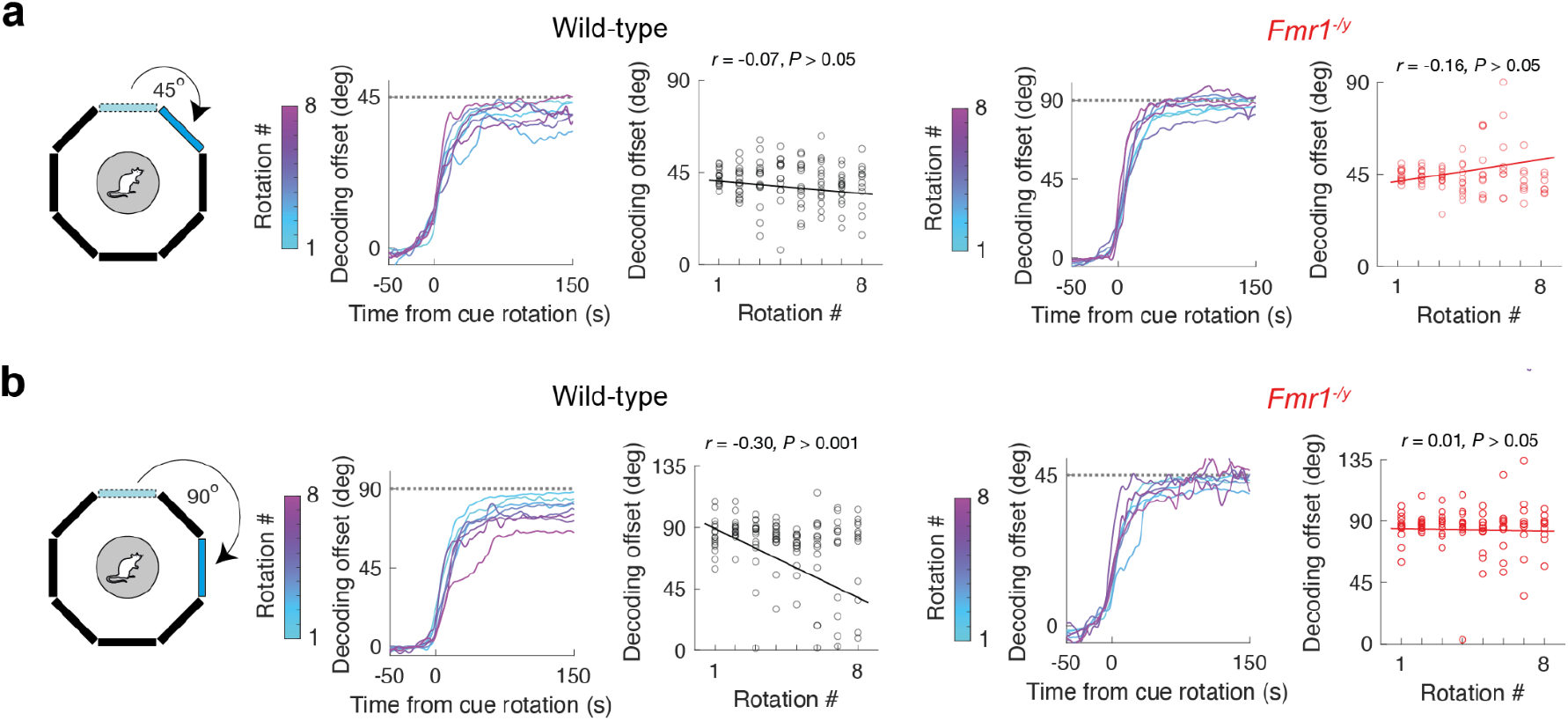
Experience-dependent reduction in the magnitude of HD signal reorientation is absent in *Fmr1*^*-/y*^ rats. **a–b**, Trial-by-trial change in HD offset measured immediately following cue rotation for WT and *Fmr1*^*-/y*^ rats. **a**, 45° cue rotations (low cue conflict). **b**, 90° cue rotations (high cue conflict). Cyan-magenta color map denotes trial averages, points denote individual rotation events across the eight rotations per session; black lines indicate the linear fit. In WT rats, reorientation magnitude did not vary systematically across repeated 45° rotations (**a**; r = -0.07, n = 128 rotations, *P* > 0.05), but progressively decreased across repeated 90° rotations (**b**; r = -0.30, n = 128 rotations, *P* < 0.001), consistent with experience-dependent down-weighting of the visual landmark under high cue conflict. In contrast, *Fmr1*^*-/y*^ rats showed no experience-dependent reduction in reorientation magnitude for either 45° rotations (**a**; r = 0.16, n = 96 rotations, *P* > 0.05) or 90° rotations (**b**; r = 0.01, n = 96 rotations, *P* = > 0.05).

We hypothesized that the reduced contribution of self-motion information to sensory integration in *Fmr1*^*-/y*^ rats may reflect diminished signalling of angular head velocity (AHV) within PoSub, thereby weakening idiothetic constraints on reorientation during cue conflict. Consistent with prior work (Spalla et al., 2022), PoSub neurons exhibited widespread AHV modulation in both putative (EX cells, often in conjunction with HD tuning (Figure 3a–b), and in fast-spiking (FS) cells (putative interneurons, n = 85 cells from 8 WT rats, n = 83 cells from 7 *Fmr1*^*-/y*^ rats), which were more broadly tuned to HD (Figure 3f–g) (Duszkiewicz et al., 2024; Tukker et al., 2015). To quantify the reliability of AHV coding across these cell classes, we computed an AHV score for each cell, defined as the correlation between the binned spike train and the animal’s instantaneous AHV (Keshavarzi et al., 2022). Both EX and FS cell populations in *Fmr1*^*-/y*^ rats showed a significant reduction in AHV coding relative to WT controls (Figure 3c, h). Moreover, FS cells, but not EX cells, exhibited a reduced dynamic range of AHV modulation, quantified as a decrease in AHV tuning-curve slope (Figure 3d, i). Together, these results indicate that AHV signalling in PoSub is weakened in *Fmr1*^*-/y*^ rats, providing a plausible circuit-level substrate for reduced self-motion weighting during reorientation.

**Figure 3.**
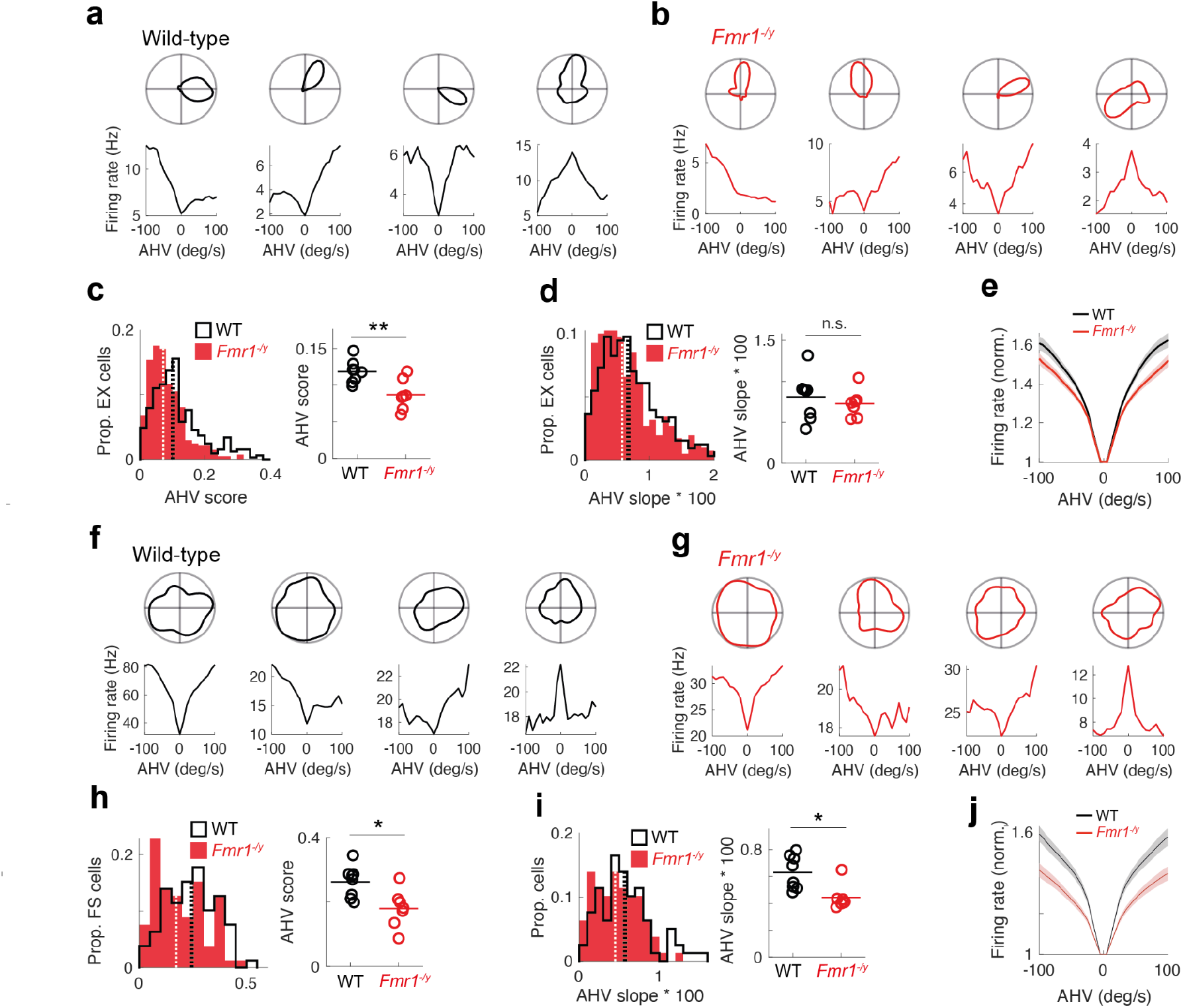
Attenuation of AHV coding across PoSub cell populations in *Fmr1*^*-/y*^ rats. **a-b**, examples of HD tuning curves (top) and AHV tuning curves (bottom) of four EX cells in WT rats (**a**) and Fmr1^-/y^ rats (**b**), showing various AHV tuning profiles. **c-e**, EX cells in *Fmr1*^*-/y*^ rats show degraded AHV tuning in comparison with WT rats (**c**, AHV score, LME: genotype effect, *P* < 0.01) but exhibit WT-like dynamic range in their AHV tuning curves (**d-e**, AHV slope, LME: no genotype effect, *P* > 0.05). **f-g**, examples of HD tuning curves (top) and AHV tuning curves (bottom) of four FS cells in WT rats (**f**) and Fmr1^-/y^ rats (**g**), showing various AHV tuning profiles. **h-j**, FS cells in *Fmr1*^*-/y*^ rats show degraded AHV tuning (**h**, AHV score, LME: genotype effect, *P* < 0.05) and attenuated dynamic range in their AHV tuning curves (**i-j**, AHV slope, LME: genotype effect, *P* < 0.05) in comparison with WT rats. Dashed lines on the histograms indicate population medians; horizontal lines on animal average panels are means of animal averages; shaded areas indicate +/-SEM.

## Discussion

In the present study, we leveraged the cortical HD circuit to examine the mechanisms of sensory integration of visual landmark and self-motion information in a rat model of Fragile X Syndrome. Following previous reports on a single-cell level (Rotenberg and Muller, 1997; Zugaro et al., 2000; Knight et al., 2014), we confirmed that in WT rats, rotations of a visual landmark in the presence of the animal induced only partial shifts in the anchoring of individual tuning curves as well as the population-level HD signal, reflecting a weighted integration of these two sources of orientation information. In *Fmr1*^*-/y*^ rats, however, the reorientation events were fully dominated by the visual cue: reorientation of the HD signal precisely followed both 45° and 90° rotations. Importantly, these differences were accompanied by a reduction in the degree of AHV tuning in both EX and FS neurons in the PoSub of *Fmr1*^*-/y*^ rats, as well as a reduction in AHV tuning slope specifically in PoSub FS cells, suggesting that weakened self-motion signalling may contribute to increased landmark capture. Collectively, our findings show that a loss-of-function mutation in a gene associated with ASD/ID in humans shifts the balance of inputs to the cortical HD circuit during cue conflict, favouring visual landmark control over idiothetic constraints.

To assess the nature of the landmark/self-motion conflict, we analysed repeated reorientation events and found that, in wild-type rats, the shifts became progressively smaller in repeated 90° rotations. In these instances, the mismatch between the visual landmark and the animal’s sense of self-motion information is higher than that of 45° rotations (Rotenberg and Muller, 1997; Knight et al., 2014). The weakening of the visual landmark control over the HD cell system with repeated shifts of the landmark in the animal’s presence suggests that the animal’s sense of self-motion becomes more salient and is consistent with an experience-dependent updating of landmark reliability (Jeffery et al., 2016; Page and Jeffery, 2018). In *Fmr1*^*-/y*^ rats, the shifts in the orientation of the HD cells remained stable across repeated conflicts, with minimal evidence for trial-by-trial recalibration in the 90° rotations. This is consistent with a diminished self-motion input to the HD cell system in adult *Fmr1*^*-/y*^ rats.

These findings raise the question of the origin of the visual landmark and self-motion signals integrated during reorientation, and how these sources might be differentially impacted in *Fmr1*^*-/y*^ rats. PoSub is a convergence point between a vestibular-driven self-motion stream (Taube, 2007) and landmark-related visual inputs conveyed through cortical and/or thalamocortical pathways (Yoder et al., 2011b). Idiothetic self-motion information is thought to enter the subcortical HD circuit through brainstem vestibular nuclei and associated pathways that compute AHV and use it to update the internal estimate of HD in the dorsal tegmental nucleus and the lateral mammillary nucleus. This pathway offers a mechanistic substrate for the internal continuity of the HD signal and a possible origin of the AHV-correlated modulation in PoSub. In parallel, multiple cortical routes could deliver allothetic visual evidence to PoSub (Street and Jeffery, 2024). Candidate sources include direct projections from the visual cortex as well as indirect cortical inputs via the laterodorsal nucleus of the thalamus (LD) (Li et al., 2025) or retrosplenial cortex (RSC), which receives rich visual information and contains cue-locked representations that are well positioned to convey landmark anchoring signals (Jacob et al., 2017; Sit and Goard, 2023). Importantly, prior causal work suggests that PoSub is not merely a passive recipient of an already-anchored HD signal: PoSub lesions disrupt landmark control of HD cell activity upstream, consistent with the idea that PoSub participates in stabilizing the mapping between the internal HD estimate and external visual reference frames (Goodridge and Taube, 1997). Thus, the heightened landmark capture we observed in *Fmr1*^*-/y*^ rats could reflect an altered balance between these two sensory streams in PoSub, shifting the circuit toward a regime in which visual inputs dominate cue conflict resolution. Establishing whether this is caused by altered AHV tuning at the level of brainstem or by weakened transmission of AHV signals to the cortex is an important future research avenue.

The current results complement recent findings of alterations in the functional development of the HD system in *Fmr1*^*-/y*^ rats (Moore et al., 2025). In this previous study, enhanced HD tuning was observed in juvenile *Fmr1*^*-/y*^ rats compared with WT littermates and reduced HD signal stability was observed in adult *Fmr1*^*-/y*^ rats (again relative to WT littermates). This instability may relate to the salience of the visual landmark. In the previous work, a small card affixed to the wall of a cylindrical recording enclosure served as the polarising landmark. If, as the current results suggest, *Fmr1*^*-/y*^ rats have a diminished representation of self-motion, instability may arise when this system is needed - for example, when the sole landmark in the environment is small and not readily visible from all orientations. In the current study, a large, illuminated LED panel was used as both the polarising landmark in the environment and its sole light source. As such, it may have been readily perceived from all directions in which the rat may face. Definitive support of this explanation, however, will require parametric variation of landmark salience in a cue-controlled environment.

More broadly, atypical sensory integration is increasingly viewed as a prominent circuit-level phenotype in ASD/ID (Meilleur et al., 2020; Siemann et al., 2020; Wallace et al., 2020). This may reflect altered sensory coding within individual modalities (Falcão et al., 2024; Mansour et al., 2021; Marco et al., 2011; Williams et al., 2023) and/or disrupted binding of unimodal signals into coherent multisensory percepts within higher-order association cortices. In both humans (Gao et al., 2023) and rodents (Garner and Keller, 2022; Meijer et al., 2018), multisensory integration has most often been studied using audiovisual paradigms, and meta-analytic evidence indicates that audiovisual integration is frequently reduced in ASD (Feldman et al., 2018), with individual differences in sensory integration associated with symptom severity. At the same time, sensory features in ASD are notably heterogeneous: hyper- and hypo-sensitivity can vary independently across modalities, and sensory profiles are often not well captured by a single global dimension (Williams et al., 2023). This heterogeneity argues against a unitary explanation in terms of generalized hyper- or hypoexcitability. Instead, altered sensory integration in ASD/ID may arise from disrupted gain control of particular sensory channels or modality-specific changes in the precision of sensory representations.

The current findings provide evidence of a modality-selective decrease in sensory gain in the HD cell system of *Fmr1*^*-/y*^ rats that may relate to altered multisensory integration. These results suggest that altered sensory integration in ASD/ID may manifest not only as impaired multisensory binding per se, but also as a shift in the relative influence of specific sensory channels on ongoing cortical computations, for example by enhanced reliance on external sensory inputs when vestibular-derived self-motion signals are weakened or underrepresented.

## Methods

### Animals

All experimental procedures were performed under a UK Home Office Project Licence, approved by the University of Edinburgh Animal Welfare and Ethical Review Board (AWERB), and were conducted in accordance with the UK Animals (Scientific Procedures) Act 1986. Long–Evans male *Fmr1*^*-/y*^ rats and wild-type (WT) littermates were bred in-house by crossing WT sires with *Fmr1*-heterozygous dams. The original nul l line was generated using zinc-finger nucleases targeting *Fmr1* exon 1 with a construct containing the eGFP coding sequence, as previously described (Asiminas et al., 2019). Males from this line do not express either FMRP or eGFP.

For the present experiments, rats were reared with parents and littermates until weaning (P21), after which they were housed in mixed-genotype littermate cages. Animals were maintained on a 12 h light/dark cycle (lights on at 07:00) and provided standard laboratory chow *ad libitum* prior to behavioural training. Before habituation, rats were placed on mild food restriction to encourage foraging, with body weight maintained at ≥85% of free-feeding weight. Animals were returned to *ad libitum* feeding 48 h prior to surgery and received supplementary water-soaked breeding chow during the post-operative period. Mild food restriction was reinstated 24 h after surgery ahead of the experimental protocol.

Rats were randomly selected from litters for inclusion in the study. Experimenters remained blinded to genotype throughout surgery, recordings, and preprocessing, and were unblinded only at the analysis stage.

### Surgery

Rats were anaesthetised using 1.5-3.5% isoflurane, while the animal’s body temperature was maintained at 37^°^C using a homeothermic blanket system (Harvard Apparatus). Carprofen (Rimadyl, 50 mg/mL) was administered subcutaneously at 0.08 mL/kg body weight shortly after induction of anaesthesia. To secure the headcap and implant on the head, the skull was covered with dental adhesive (Optibond Universal). This was supplemented with three to five self-tapping jeweller’s screws. A silver ground wire was attached to one of the skull screws placed above the cerebellum in adult rats. A craniotomy was drilled over the left postsubiculum, and a silicon probe (Cambridge Neurotech, model H7) was lowered into the brain, slightly above the target area (adults, from Bregma: AP, −7.64 mm; ML, 3.30 mm; DV, −1.70 mm). The silicon probe was mounted on a drive (3DNeuro or made in-house), secured onto the skull’s surface using dental acrylic. Animals were also implanted with an additional probe placed in the retrosplenial cortex. After surgery, the animals were allowed to recover on a heated bench (30°C) for at least one hour and returned to the animal house for at least 24 hours before recording. After surgery rats were either single-housed or housed with a sibling in a cage with a divider separating them. Adult rats were at least 8 weeks old at the time of implantation.

### Recording

The probe was initially implanted in the white matter above PoSub and was subsequently advanced slowly into PoSub (speed: 0.28 mm/h in 0.035 mm increments) 24h after implantation. The tissue was allowed to settle for at least 2h before recordings commenced.

The recording rig consisted of an aluminium frame (138 cm x 138 cm x 188 cm height) with aluminium floor, ceiling and walls (60 cm height), with matt black vinyl flooring covering the floor of the frame as well as the inner walls. The frame was painted with a matt finish black barbecue paint (Hammerite) and was surrounded by black curtains to minimise the influence of uncontrolled distal cues. The exploration platform (50 cm diameter, 2 cm lip) was placed in the centre of the frame and was elevated 16 cm above the floor of the frame. A pellet dispenser (CI.80209A-45, Campden Instruments) was mounted on top of the frame to deliver sugar pellets through a tube protruding through the hole in the centre of the ceiling. Inside the enclosure there were 8 LED screens (made in-house; 50 cm width, 50 cm height, 2.5 cm thickness), arranged in an octagonal shape. Each screen consisted of three 16x16 RGB LED panels (Adafruit Neopixel WS2812B, 1cm LED spacing) arranged horizontally so that the bottom row of LEDs was 26 cm above the floor level, and placed within a casing made of two perforated aluminium sheets (RS, 50 cm x 50 cm, hole diameter: 2mm, open area percentage: 47%) to hide the LED panels. The sheets were painted with matt finish black barbecue paint (Hammerite) and the casing caused them to appear black when the LEDs were switched off. Each LED screen was controlled with a Raspberry Pi Pico running Adafruit CircuitPython NeoPixel library, and the individual LED screens were synchronized using a Raspberry Pi running custom Python code. The video was recorded through an acA1300-75gc Basler camera with a LMVZ4411 Kowa lens, positioned on the ceiling of the enclosure. Four speakers were placed in the centre on top of the ceiling facing radially and white noise was played through them for the duration of the recording.

Before surgery, rats were habituated to two open-field environments (a cylinder and a square arena) over three consecutive days. On recording days, animals were brought into the curtained recording enclosure and connected to the recording cable. They were then placed in an opaque bucket containing bedding and covered with a lid for 90 min, during which sleep activity was recorded. Immediately prior to the cue-rotation protocol, a masking odour (air freshener) was applied to the floor of the enclosure and the room lights were switched off. In darkness, the experimenter transferred the rat from the bucket onto the elevated platform, at which point a single visual cue was displayed on one LED screen. The experimenter then exited the room and monitored the animal via live video feed from the adjacent control room. The cue-rotation session consisted of three phases. A baseline period (600 s) was first recorded with the cue held at a fixed reference location (0°). The cue was then displaced to an adjacent screen (45°) and subsequently returned to the 0° location. This sequence was repeated using the screen adjacent to the 45° location (90°). Each rotation was followed by a 200 s hold period during which the cue remained stationary. In total, each session contained four blocks of four cue rotations (16 rotations total: eight clockwise and eight counterclockwise). To sustain locomotion across the full protocol, sugar pellets were delivered intermittently from an overhead feeder.

To reduce the influence of uncontrolled auditory cues, white noise was continuously played through the overhead speakers throughout the session. The arena was swept and cleaned with chlorhexidine solution (Vetasept) before and after each recording session. Each rat underwent the protocol twice, using a different elevated platform (green or blue), visual cue pattern (blue crosshatched gratings or teal vertical stripes), initial cue positions (right or bottom), direction of the initial rotation (counterclockwise or clockwise) and masking odour across sessions to maximize novelty and encourage exploration.

Electrophysiological signals were recorded through a headstage (RHD, Intan technologies) connected to an Intan RHD standard SPI cable, which was connected to a custom-made commutator in the ceiling. Signal was acquired at 30 kHz with an Open Ephys data acquisition system. For tracking the rat’s position and head direction, a red and a green LED were attached on the sides of the headstage. Video capturing and LED tracking were performed using Bonsai software (Lopes et al., 2015) at 40 Hz. The camera sent a TTL pulse to the data acquisition system at the time of each frame, allowing the synchronisation of the two streams of information.

### Histology

After data collection the rats were anaesthetised with 5% isoflurane and injected intraperitoneally with a lethal dose of pentobarbitone (≥1ml for adults and ≥ 0.3ml for juvenile rats). After their reflexes were gone, they underwent transcardial perfusion first with 0.9% saline, and then with 4% formalin or paraformaldehyde solution. After perfusion, the brains were stored in 4% formalin or paraformaldehyde solution for at least 48 h, and then cryo-protected in a 30% sucrose solution for 48-72 h. Brains were then sliced in 40 μm coronal sections on a cryostat microtome, stained using a 0.1% cresyl violet solution and imaged with a MicroPublisher 3.3 RTV camera (Qimaging) on a Northern Light precision illuminator (Imaging Research Inc), using Image-Pro Premier 9.2 software.

### Spike sorting and cell classification

Spikes were first split automatically into clusters using Kilosort 2.0 (Pachitariu et al., 2016). Manual curation was performed while visualising the automatically created clusters on Klusters software. At this stage, any cluster without a clear waveform and clear dip in the spike train autocorrelogram at the 0-1 ms time bin was classified as noise and cluster pairs with similar waveforms and a clear dip in their spike train cross-correlograms at the 0-1 ms time bin were merged. Viable units were first identified as units that had a waveform with negative deflection (criterion aiming to exclude spikes from fibers of passage). Next, a cluster contamination score was computed from the spike autocorrelograms, as a ratio of the amplitudes of the 0-1 ms bin and the maximum amplitude within 0-100 ms. Units with cluster contamination scores above 0.5 were excluded from the analysis. Next, putative excitatory cells were classified on the basis of trough-to-peak duration of their average waveforms. The classification threshold was set to 0.45 ms, corresponding to the trough of the bimodal distribution of trough-to-peak durations. For each analysis, units with positive waveforms and units with less than 100 spikes fired in the analyzed epoch were excluded.

### HD tuning curves

The animal’s HD was calculated based on the line perpendicular to the line connecting the red and green LEDs attached to the headcap. HD tuning curves were then computed as the mean firing rate of the cell in each 1^°^ angular bin and smoothened with a circular Gaussian window (6^°^ SD). Speed was computed as the difference in the animal’s position at successive time points and was smoothed with a moving average window of 0.25 s. HD tuning curves were computed from epochs when the animal’s speed exceeded 1.5 cm/s. HD information was calculated from tuning curves containing N angular bins as follows:

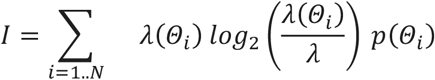

where λ(Θ_*i*_) is the firing rate of the cell for the *i*th angular bin and λ is the average firing rate of the neuron during exploration. We obtained HD tuning curve crosscorrelograms by computing Pearson’s correlation coefficients between the reference tuning curve vector and the second tuning curve vector (from either the same or another cell), which was circularly shifted by 1 to maximum number of bins.

### Bayesian decoding and drift estimation

To estimate the degree of drift of the internal HD with respect to the allocentric reference frame, spike times of individual cells were binned into population vectors (200 ms window, smoothed with a 400 ms SD Gaussian window). Since most cells in PoSub show some degree of HD tuning (Duszkiewicz et al., 2024), we used all cells that fired more than 100 spikes in a given epoch to construct the population vectors. Based on cells’ tuning curves from the baseline period (i.e. period before the first cue rotation), the population vectors were converted into a Bayesian probabilistic map under the hypothesis that HD cells fire as a Poisson process (Zhang et al., 1998). The instantaneous internal HDs were taken as the maxima of these probabilistic maps, and the values corresponding to the probabilities below 1SD of the probability maxima distribution were discarded. The degree of drift of the HD signal was calculated as the decoder error: angular difference between the real HD and the decoded HD at each time bin. To remove the noise fluctuations in the resulting drift signal, decoder error was denoised with a hampel filter (5 s window, 1 SD) and then smoothened with a 0.8 s SD Gaussian window.

To compute the change in offset around the time of the cue rotation, the drift values around the cue rotation (200 s intervals, from 50 s before to 150 s after each cue-rotation event) were normalized by subtracting the median drift value over 50 s before the cue rotation event. The values around counterclockwise rotations were then multiplied by -1 to match them to the clockwise rotations. Resulting offset traces were then included in further analyses according to the following analysis when (1) the median offset after cue rotation was higher than the median offset before cue rotation, (2) the circular variance in the offset before or after cue rotation was below 20. The post-rotation offset values were then computed as the median value from 30 s to 150 s after each cue rotation.

### Angular velocity tuning

Angular head velocity (AHV) was calculated as the angular difference between HD values at successive time points and was smoothened with a Gaussian window (0.25 s SD). AHV tuning curves were computed as a mean firing rate of a cell in each AHV bin (10 deg/s bin size) up to the maximum of 100 deg/s and smoothened with a Gaussian window (20 deg/s SD). The slope of the linear fit was then computed separately for each side of the tuning curve (clockwise and counterclockwise) and the higher absolute slope value out of the two was used for comparisons. For AHV score, we first defined separate epochs of clockwise and counterclockwise AHV (minimal angular speed of 2 deg/s). Binned spike times of each cell (200 ms window, smoothed with a 400 ms SD Gaussian window) were then correlated with the AHV values at each time bin, separately for the clockwise and counterclockwise epochs, and the higher absolute value out of the two correlation coefficients was used as the AHV score.

### Data analysis and statistics

All analyses were conducted using Matlab R2020b (Mathworks) with TStoolbox (https://github.com/PeyracheLab/TStoolbox) and Circular Statistics Toolbox version 1.2167 (Berens, 2009). Statistical comparisons of individual cell data were performed using a generalized linear mixed model (GLMM) approach (Yu et al., 2022) in order to take into account the hierarchy of dependency in our data sets (genotypes-rats-neurons) and account for random effects. Statistical modelling routines for the linear mixed effects (LME) models were written and run using RStudio 2023.12 (RStudio Team). Depending on the data distribution (normal, lognormal or gamma), linear mixed models (LMMs) or generalized linear mixed models (GLMMs) were fitted to single unit data metrics using the R package lme4 v1.1–17. Animal and cell identity were included in models as random effects, and genotypes were included as fixed effects. Terms are progressively eliminated when a simpler model (i.e. a model not containing that term) fits the data equally well (based on likelihood ratio test). Consequently, the p-values reported in the context of LMEs are given by likelihood ratio tests between a model containing the variable or interaction in question and a model without that variable or interaction (a reduced/null model). All statistically significant differences obtained through LMEs were also statistically significant when comparisons were made using a two-sample Mann-Whitney U test with cells as experimental units.

Statistical comparisons on animal averages (where stated) were performed with independent or paired Student’s t-tests. All statistical tests were two-tailed. No statistical methods were used to predetermine sample sizes, but our sample sizes are similar to those reported in previous publications.

## Data Availability

Data will be made openly available at the time of publication.

## Code Availability

Code will be made openly available at the time of publication.

## Acknowledgements

This work was supported by the Simons Foundation Autism Research Initiative grant AN-AR-Rat Models-0090333 (AJD, AV, PAD, and AP), the Canadian Research Chair in Systems Neuroscience (AP), CIHR Project Grants 155957 and 180330 (AP); NSERC Discovery Grant RGPIN-2018-04600 (AP) and Simons Initiative for the Developing Brain (ERW, AR). The authors thank Lynn Morrison and the other animal technicians from the University of Edinburgh George Square Bioveterinary Services Facility for their expertise and assistance in this work, and Dr Owen Dando for help implementing the LME analysis.

## Declaration of Interests

The authors declare no competing interests in relation to this work.

**Figure S1.**
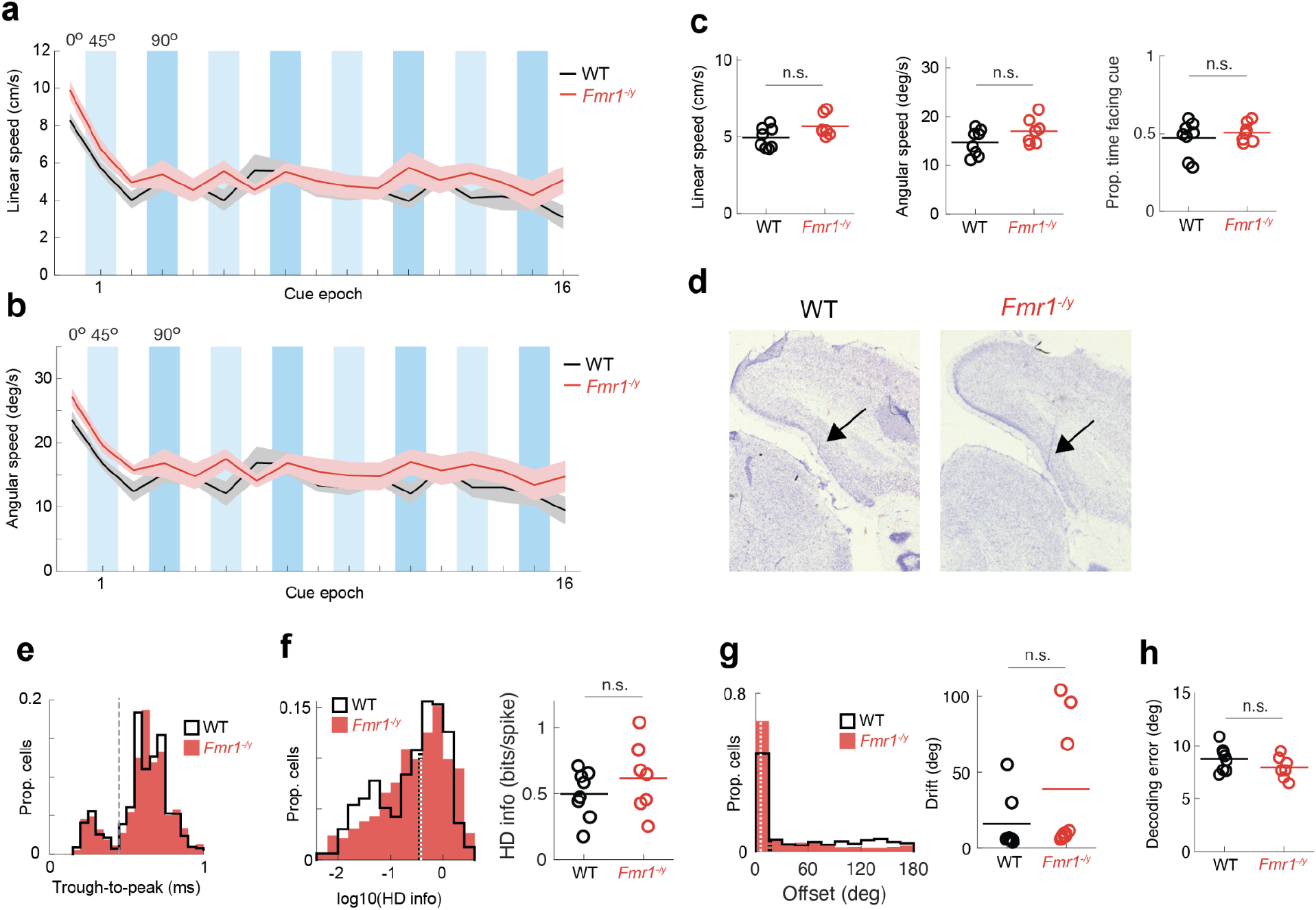
Behaviour, probe placement and basic tuning properties in WT and *Fmr1*^*-/y*^ rats. **a-b**, average linear speed (**a**) and angular speed (**b**) for each of the task epochs. **c**, No differences in the average linear speed (left), angular speed (right), or the proportion of time spent facing the cue (+/-90 degrees) across the whole task session. (Linear speed, t(13) = 2.07, *P* > 0.05; angular speed, t(13) = 1.72, P > 0.05; prop. time facing the cue, t(13) = 0.75, *P* > 0.05). **d**, Example silicon probe tracts going through PoSub, as visualized by Nissl staining. Black arrows, electrode tracts. **e**, Distribution of the trough-to-peak duration of all recorded units. Grey dashed line, threshold of cell classification into putative excitatory cells (EX) and putative fast-spiking interneurons (FS) based on the trough in the bimodal distribution. **f**, Overall distribution (left) and animal averages (right) of HD info indicate similar degree HD tuning in WT and *Fmr1*^*-/y*^ rats (LME: no genotype effect, *P* > 0.05). **g**, distribution of tuning curve offsets between the initial baseline period and the end period of the recording session for all cells, as well as animal averages, indicating similar degree of anchoring of the HD signal to the allocentric reference frame in WT and *Fmr1*^*-/y*^ rats (angular offset between tuning curves, LME: genotype effect, *P* > 0.05). **h**, median absolute decoding error computed between two halves of the initial baseline epoch indicates similar decoding quality across WT and *Fmr1*^*-/y*^ rats. Dashed lines on histograms, population medians; horizontal lines on animal average panels, means of animal averages.

